# Long dsRNA on the move: Extracellular vesicles deliver dsRNA for antiviral protection in human cells

**DOI:** 10.64898/2026.07.09.737430

**Authors:** Dominique E. Daniels, Shawn M.S. Carr, Stephanie J. DeWitte-Orr

**Affiliations:** Department of Biology, Wilfrid Laurier University, Waterloo, ON, Canada; Department of Health Sciences, Wilfrid Laurier University, Waterloo, ON, Canada

**Keywords:** Extracellular vesicles, long double stranded RNA, human coronavirus 229E, vesicular stomatitis virus, U937, HEL-299, type 1 interferon

## Abstract

Viruses make long (>40 bp) double stranded RNA (LdsRNA) during replication, which stimulates the innate antiviral immune response. In vertebrates, LdsRNA can induce the type I interferon response (IFN) or the antiviral RNA interference response (dsRNAi) to limit viral replication. Extracellular vesicles (EVs) have previously been shown to carry a variety of nucleic acids for intercellular signaling, and insects and plants have been shown to package LdsRNA in EVs as part of their antiviral immune response. We hypothesized that a similar phenomenon occurs in vertebrates in which EVs traffic LdsRNA between cells during viral infection to induce an antiviral response in naïve cells. In this study we showed that both vesicular stomatitus virus (VSV)-derived and *in vitro* transcribed (ivt)-LdsRNA can be packaged into EVs. LdsRNA was detectable by immunoblots in EVs extracted by both differential ultracentrifugation from ivt-LdsRNA treated U937 and ExoQuick-TC^TM^ precipitation from VSV-infected U937 cells (dsRNA-EVs) but not uninfected controls (control EVs). Isolated EVs were roughly 100 nm in diameter and were able to protect LdsRNA from degradation by RNase III. LdsRNA delivery by dsRNA-EVs was visualized in HEL-299 cells via immunocytochemistry (ICC). The LdsRNA-EVs protected against infection from HCoV-229E, while control EVs did not. Together these results indicate EVs can package and deliver long dsRNA to provide antiviral protection in naïve vertebrate cells.

**Author Summary:** When viruses infect cells, they produce double-stranded RNA, a molecule that alerts the body to the presence of infection and triggers antiviral defenses. Previous studies have shown that cells can release small membrane-bound packages called extracellular vesicles, which carry biological messages to other cells. However, it was not known whether antiviral double-stranded RNA could be transported in these vesicles and shared with neighboring cells.

In this study, we investigated whether human cells package double-stranded RNA into extracellular vesicles and whether this cargo helps protect other cells from viral infection. We found that both synthetic and virus-derived double-stranded RNA were incorporated into extracellular vesicles and shielded from degradation. These vesicles successfully delivered double-stranded RNA to untreated cells, substantially protecting these cells from infection with a human coronavirus. Our findings suggest that cells can communicate antiviral warnings to neighboring cells by packaging double-stranded RNA into extracellular vesicles.

This work reveals a previously unrecognized way that antiviral protection may spread through tissues during infection. By extending antiviral signals beyond directly infected cells, extracellular vesicles may help coordinate a broader host defense response. Understanding this natural communication system could also inform the development of new RNA-based antiviral therapies.

## Introduction

Long (>40 bp) double stranded RNA (LdsRNA), is made by viruses during their replication cycle (1,2). In vertebrate cells, high concentrations of LdsRNA are sensed as a pathogen associated molecular pattern (PAMP) by the cell’s pattern recognition receptors (PRRs), including toll-like receptors (TLRs), and retinoic acid-inducible gene I (RIG-I) like receptors (RLRs) triggering the type I interferon (IFN) response, (reviewed by 3,4). Alternatively, at low concentrations, LdsRNA can be cleaved by the endonuclease Dicer, activating the RNA interference (RNAi) pathway, called the dsRNAi pathway (5). Extracellular dsRNA can enter the cell via class A scavenger receptors (SR-As) through clathrin mediated endocytosis (6). Once in the endosome, the dsRNA can be sensed by TLR3 or escape to the cytoplasm via the SIDT2 channel where it can be detected by RIG-I, melanoma differentiation-associated protein 5 (MDA-5) or Dicer. Detection by TLR3, RIG-I or MDA-5 results in the expression of IFNs such as IFN-β, which promotes the expression of IFN stimulated genes (ISGs) via the Janus kinase (JAK)-signal transducer and activator of transcription (STAT) pathway. ISGs inhibit viruses throughout their replication cycle including virus entry (e.g., CH25H, IFITM family), protein production and modification and viral egress (e.g., PKR, ISG15) (7–11). Detection by Dicer results in viral LdsRNA cleavage into small interfering (si)RNAs which when loaded into the RNA-induced silencing complex (RISC), suppresses translation of viral mRNAs whose sequences match the siRNAs produced inhibiting viral protein synthesis and subsequent viral replication (Reviewed by (12,13).

Extracellular vesicles are small lipid bound particles that are released from cells for a variety of physiological functions from intercellular communication to waste elimination. A wide array of RNA species are packaged into EVs including mRNAs, long non-coding (lnc)RNAs, PIWI-interacting (pi)RNAs, transfer (t)RNAs and small nuclear (sn)RNAs (14–16). Recent studies demonstrate that small RNAs and synthetic polyinosinic:polycytidylic acid (poly IC) can be incorporated into EVs in a rheumatoid arthritis model (17–19). While there is evidence that HSV-1 derived LdsRNA has been detected in and shown to stimulate an innate immune response in uninfected neighbouring cells, whether virus-derived long dsRNA is actively packaged into EVs in vertebrate systems and whether such EV-associated dsRNA can functionally induce antiviral protection in recipient cells remains unresolved (20). In contrast, plant systems demonstrate EV-mediated spread of both siRNAs and long dsRNA as part of systemic dsRNAi responses (21).

Given the central role of extracellular dsRNA in innate immune activation, we sought to determine whether EVs could package and deliver long dsRNA capable of inducing an antiviral response in naïve cells. To address this, we utilized U937 cells as EV-producing cells and HEL-299 cells as interferon-competent naïve recipient cells permissive to HCoV-229E infection (22). We tested whether EV-associated long dsRNA could be transferred to recipient naïve cells and assessed its impact on antiviral protection.

## Results

### Extracellular Vesicles from U937 can package *in vitro* synthesized and virus-derived LdsRNA

The first step of the study was to determine whether EVs could package LdsRNA from both intracellular and extracellular sources. For extracellular LdsRNA, EVs were isolated by ultracentrifugation from U937 cultures that were treated with 33.3 µg/mL in vitro synthesized (ivt-)LdsRNA (HCoV-229e N, 700 bp, Figure 1A) or untreated (control) cultures. LdsRNA’s ability to be packaged into EVs was tested using immunoblots with the J2 antibody on the EV preparation and cell pellets from untreated and dsRNA treated cultures (Figure 1B). LdsRNA was not detected in the cell or EV pellets from untreated control U937, but was detected in both the cell extracts and EV preparation from dsRNA treated cells. Following the 16 hr dsRNA treatment (17), the LdsRNA in the cell had degraded from its original length of 700 bp, into a smear of lengths with some fragments being less than 150 bp. Furthermore, the majority of LdsRNA packaged into the EVs was found to be <300 bp. The quantity of LdsRNA taken up by the cell appears to be substantially greater than the amount of LdsRNA that becomes packaged into EVs and the length of the LdsRNA in the EVs is a range of sizes as it produces a smear instead of single band.

**Fig 1.**
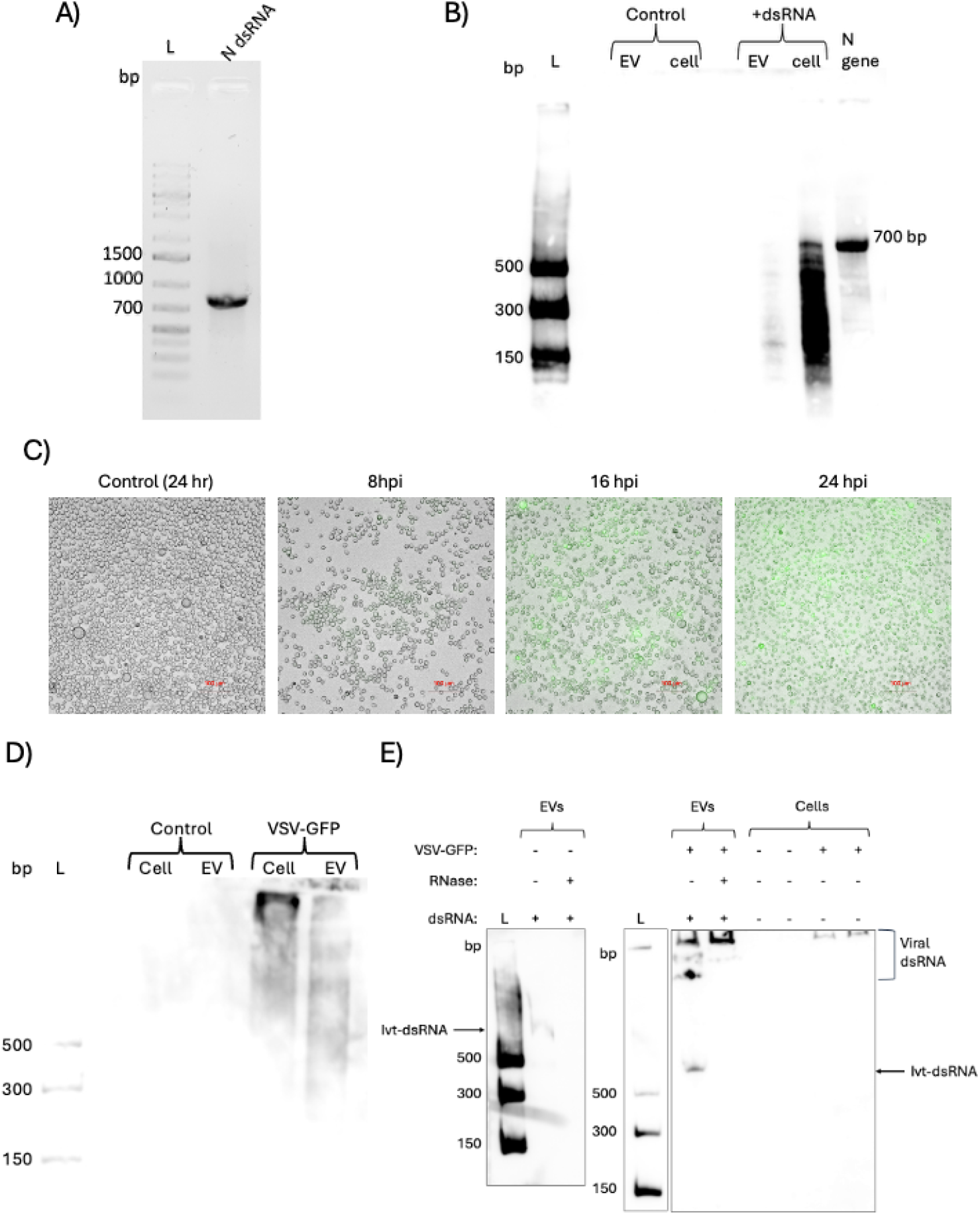
In vitro synthesized (ivt-) and virus-derived LdsRNA encapsulated in extracellular vesicles. (A) Agarose gel electrophoresis of 100 ng ivt-LdsRNA (700 bp) encoding the HCoV-229E N gene (N dsRNA). RNA was resolved on a 1% agarose gel and visualized using RedSafe. Lane L: GeneRuler 1 kb Plus DNA Ladder. (B) J2 immunoblot detection of dsRNA extracted from extracellular vesicles (EVs) and whole-cell lysates of U937 cells treated with or without ivt-LdsRNA. EVs were isolated by ultracentrifugation 16 hr following treatment with 33.3 µg/mL long dsRNA (HCoV-229E N, 700 bp). Lane L: dsRNA ladder. C) GFP production by VSV-GFP (MOI I) at 8, 16, and 24 hpi in U937 cells. Images taken at 200x magnification. D) EVs were extracted from U937 cells using ExoQuick-TC™ following infection with VSV-GFP (MOI 1) for 8 hr. Long dsRNA was detected only in infected cells and resulting EV pellets via J2 immunoblot. Lane L: dsRNA ladder. E) Cleared media was collected from VSV-GFP infected (MOI 1) U937 at 8hpi and spiked with 1µg LdsRNA. EVs were pelleted and treated with 2U RNase III prior to RNA extraction. J2 immunoblots showed that most virus-derived RNA in EVs was protected from RNAse III degradation, while spiked in LdsRNA was not. Lane L: dsRNA ladder. Data are representative of at least three independent experiments.

Next, U937 were infected with VSV-GFP (MOI 1) to determine whether intracellular LdsRNA was packaged into EVs during viral infection. VSV-GFP was selected for this experiment due to its ease of infection visualization via GFP production (23), rapid replication cycle (24), and production of detectable levels of LdsRNA (25). First, the optimal EV collection time was assessed through a time course of VSV-GFP infection in U937 cells. VSV-GFP infected U937 cells (MOI 1) were imaged at 8, 16, and 24 hr post infection (hpi) to assess GFP production and cytopathic effect (CPE) in the form of cell death. GFP expression was observed at 8 hpi and increased at 16 hpi and 24 hpi (Figure 1C). Minimal CPE was observed at 8 hpi, with more severe CPE observed at 16 hpi and 24 hpi, therefore the 8 hpi timepoint was chosen for EV collection in downstream experiments.

EVs were extracted from uninfected and VSV-GFP infected (MOI 1) U937 culture supernatant at 8 hpi using ExoQuick-TC™, an EV precipitation reagent, with uninfected and infected U937 cell pellets being used as controls. RNA was extracted from both the cells and EVs. Immunoblots with the J2 antibody showed long dsRNA was present in both the infected cells and infected cell supernatant extracted EVs at 8 hpi (Figure 1D). No long dsRNA was detected in the uninfected control U937 cells or their EVs, confirming that the long dsRNA was virus derived. The LdsRNA isolated from the virus infected cell pellets and resulting EVs represent lengths >500bp, with some lengths likely representing the length of the VSV genome [11kb; (26)]. To confirm that the LdsRNA was encapsulated in the EVs and not simply precipitated with the EVs, LdsRNA of known length (HCoV-229E N, 700 bp) was added to cleared media from VSV-GFP infected (MOI 1) U937 prior to the addition of ExoQuick-TC™. EV pellets were treated with RNase III prior to RNA extraction. Immunoblots with the J2 antibody showed most of the virus-derived LdsRNA was protected from RNAse III degradation, while the spiked in N LdsRNA was not (Figure 1E).

Ultracentrifugation was pursued for subsequent experiments because it is a physical separation method that produces EV preparations without precipitating other cellular components. Thus, EVs isolated by ultracentrifugation were characterized through transmission electron microscopy (TEM) and Western blot analysis. TEM showed that the EVs were roughly 100 nm in diameter, placing them in the small EV (SEV) category (Figure 2A) (27). EVs were found to have a cup-shaped morphology which is typical of SEVs isolated through ultracentrifugation (28). Common EV proteins HSP70, Histone H3, and EMMPRIN (CD147) were detected in the cell and EV pellets from both dsRNA-treated and untreated U937 cells (Figure 2B) (29–31). HSP70 expression was elevated in dsRNA treated cells but levels were similar in control and dsRNA-EVs, EMMPRIN was found in high amounts in the cells and low amounts in the EVs, while Histone H3 was found at low amounts in the cells, with more in dsRNA treated cells versus controls, and high amounts in control and dsRNA-EVs. Ponceau S stain (Figure 2C) showed even protein loading across wells. The distinct Ponceau profiles between cell extracts and EV fractions suggest that the EV preparations were not contaminated with appreciable amounts of cellular proteins.

**Fig 2.**
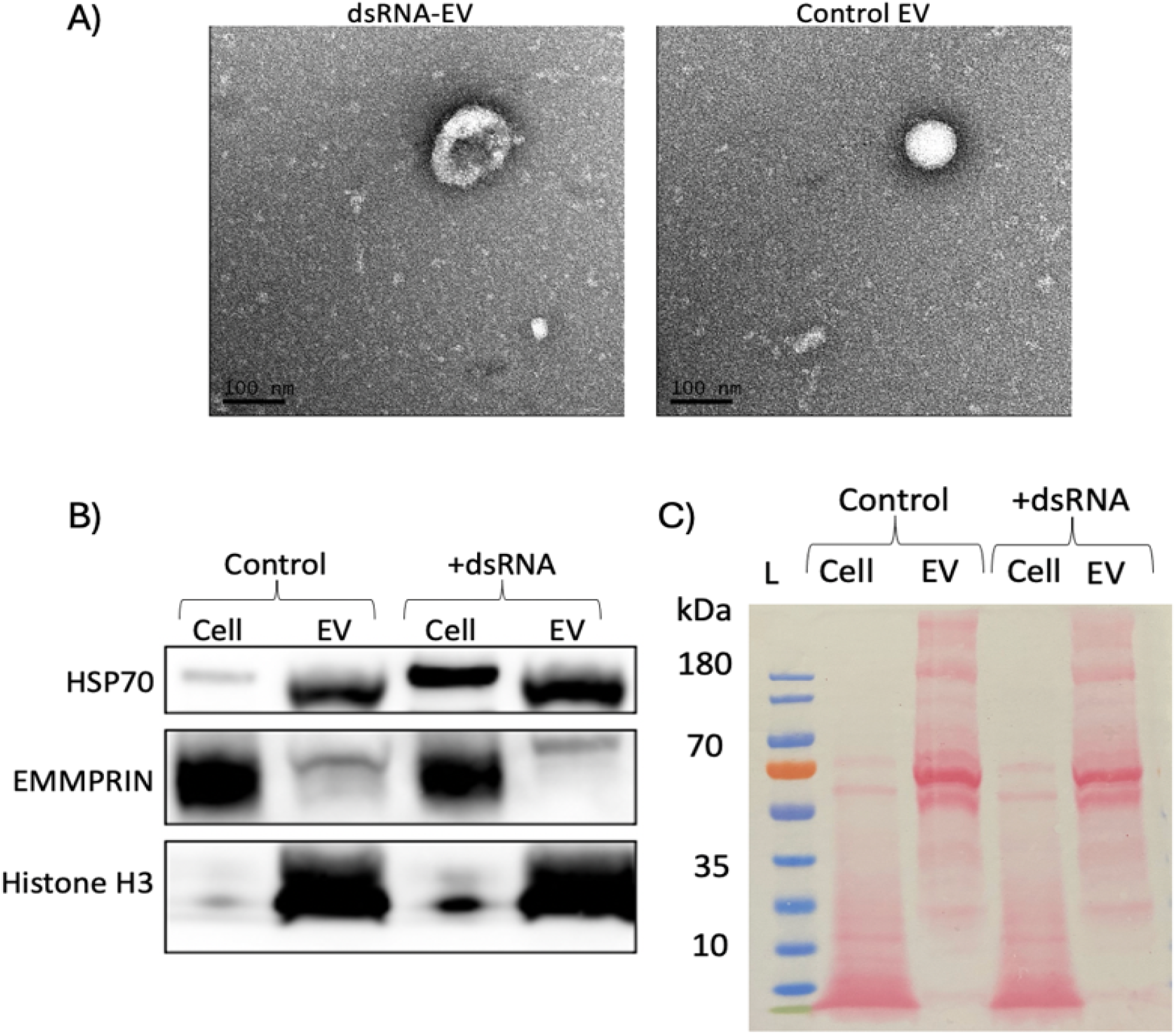
Characterization of LdsRNA-EVs. A) Transmission electron microscopy (TEM) of EVs isolated via ultracentrifugation from U937 cells treated with (dsRNA-EV) or without (control EV) 33.3 µg/mL 700 bp long dsRNA encoding the N gene of HCoV-229E. EVs were imaged at 50,000 x magnification. B) Representative Western blots showing presence of HSP70, EMMPRIN, and Histone H3 in the EV and cell pellets of both control and LdsRNA treated U937. C) Representative Ponceau S stain of total proteins on membrane prior to western blot analysis. Western blot and Ponceau S stain blots are representative of at least three independent experiments.

### Long dsRNA can be delivered to HEL-299 by Extracellular Vesicles

To determine if dsRNA-containing EVs (dsRNA-EVs) could deliver dsRNA to naïve cells, HEL-299 cells were treated with EVs isolated from LdsRNA treated (dsRNA-EVs) or untreated (control EVs) cells. After which the LdsRNA was detected by immunocytochemistry (ICC) using the J2 anti-dsRNA antibody. HEL-299 cells were selected as they are responsive to long dsRNA and have both IFN and dsRNAi pathways (5). Long dsRNA was detected in HEL-299 treated with dsRNA-EVs as well as those treated with 5 µg long dsRNA (HCoV-229E N, 700 bp) as a positive control (Figure 3). Long dsRNA was not detected in HEL-299 treated with the control EVs, which were extracted from untreated U937. Furthermore, long dsRNA was not detected in the untreated control cells. No signal was detected with secondary alone and no antibody controls.

**Fig 3.**
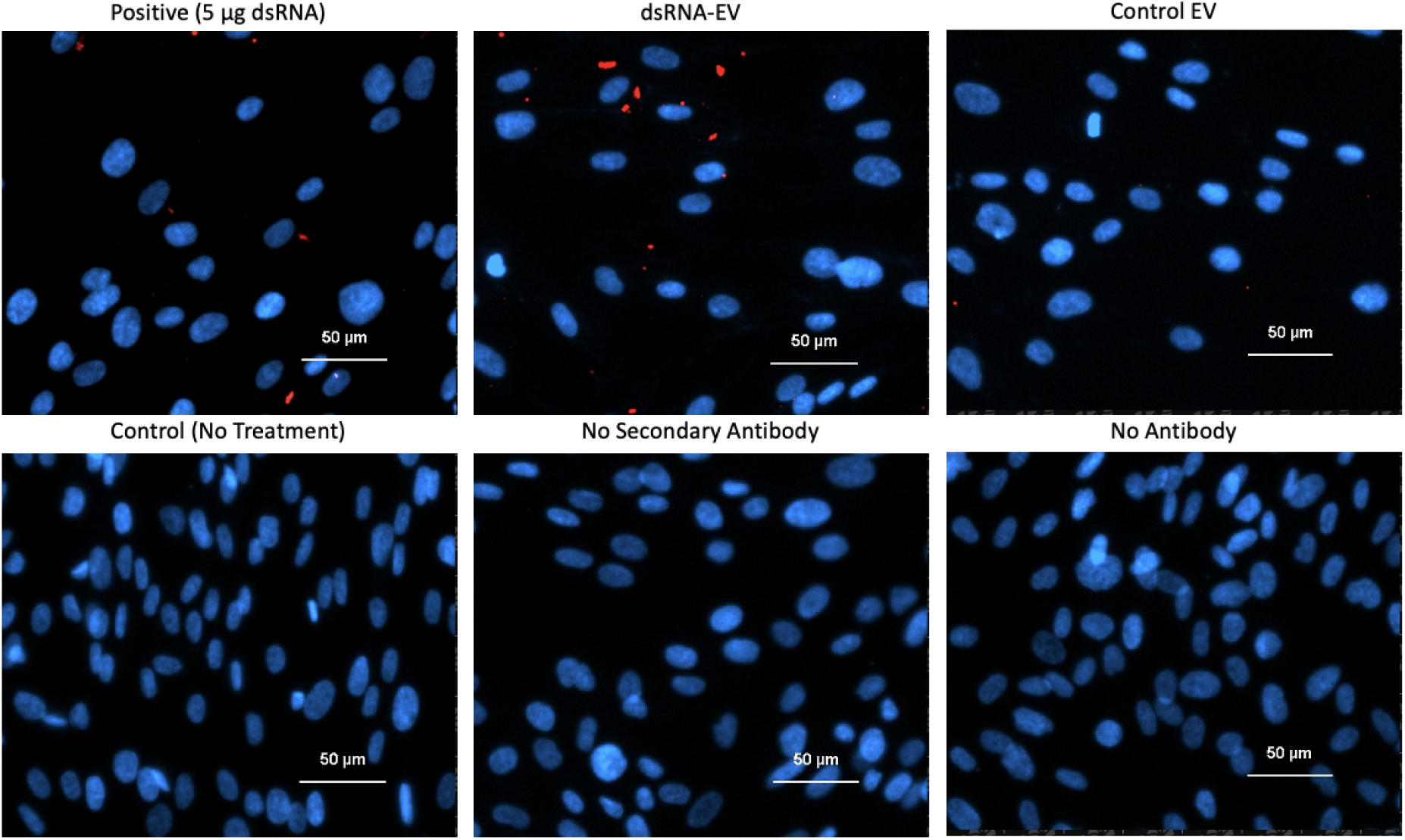
Delivery of LdsRNA to naïve HEL-299 cells by dsRNA-EVs from U937 cells. HEL-299 cells were treated with dsRNA-EVs or control-EVs isolated from U937 cultures, 5µg HCoV-229E N dsRNA (positive control), or 40µL media alone (no treatment control). Immunocytochemistry (ICC) was performed with the J2 primary antibody and goat-anti mouse TRITC secondary antibody (red). Cells were counter-stained with DAPI (blue). No antibody controls were also performed. Cells were imaged at 200 X magnification. Data are representative of at least three independent experiments.

### dsRNA-EVs provide antiviral protection against HCoV-229E infection in HEL-299

Before the antiviral activity of dsRNA-EVs could be measured, their safety as a treatment was measured. Cytotoxicity of EV treatment in HEL-299 cells was assessed via alamarBlue, and neither the dsRNA-EVs nor the control-EVs had any significant effect on cell metabolism (Figure 4A). Antiviral protection against HCoV-229E (MOI 0.065) in HEL-299 cells was then assessed following treatment with control- and dsRNA-EVs. Tissue Culture infectious Dose (TCID_50_) assays showed a significant reduction ∼2 log in viral titre in cells treated with dsRNA-EVs prior to infection (Figure 4b). Pre-treatment with control-EVs (from untreated U937 cells) provided no protection against HCoV-229E suggesting that the protection provided by dsRNA-EVs was mediated by their dsRNA cargo.

**Fig 4.**
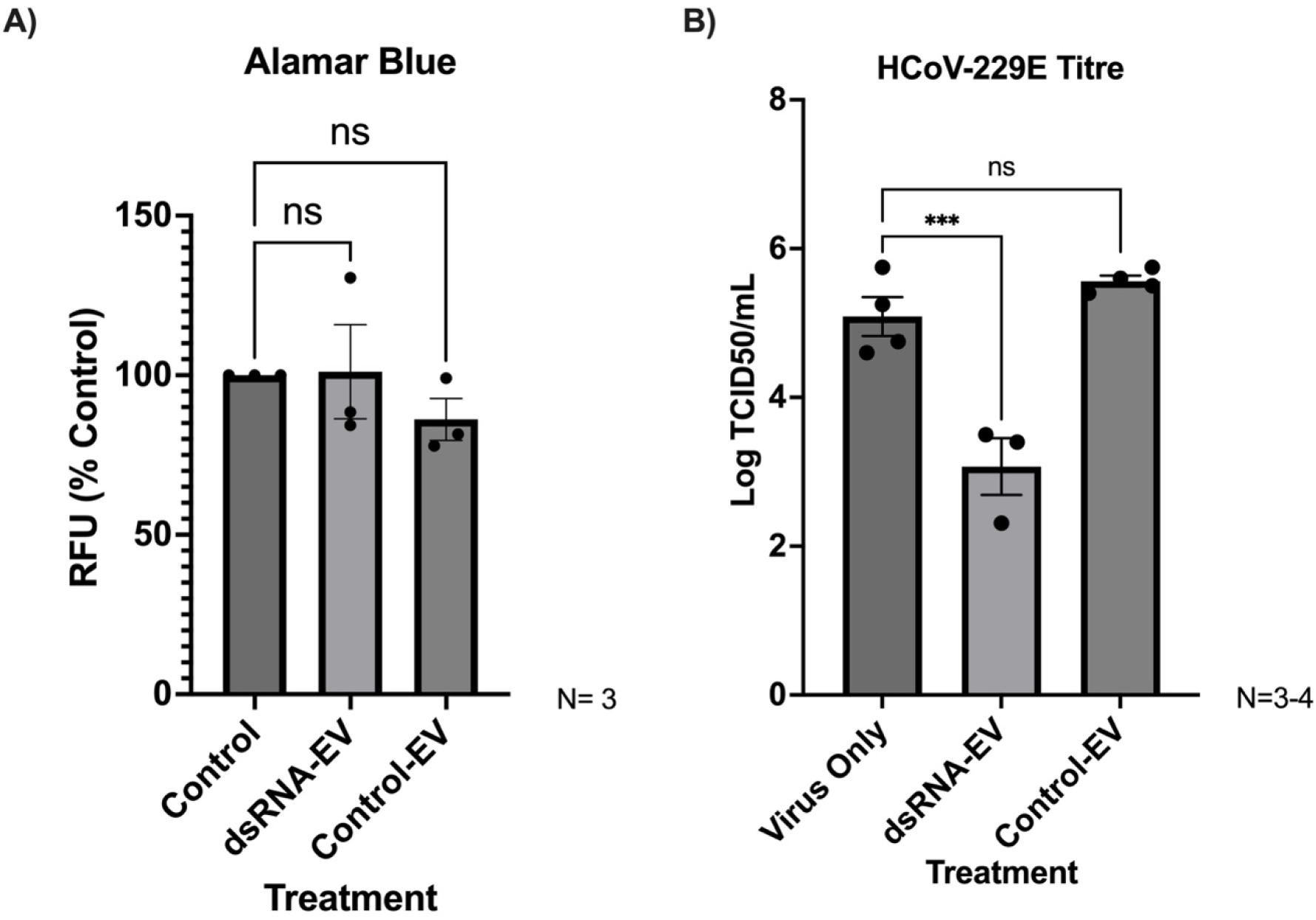
Cytotoxicity and antiviral protection from treatment with dsRNA-EVs. A) Cytotoxicity following EV treatment was measured via alamarBlue. HEL-299 were treated with dsRNA-EVs or control EVs for 24 hrs then cell viability was measured via Alamar Blue and plate reader assay. B) HEL-299 were treated with dsRNA-EVs or control EVs for 16 hrs prior to infection with HCoV-229E (MOI 0.065). Media was collected 48 hpi and virus production was quantified by TCID_50_. For both A and B: data represent the average of 3-4 independent experiments with error bars representing the SEM. Significance was determined using the Kruskal-Wallis test where a P value < 0.05 was considered significant.

## Discussion

EVs play many important roles in intercellular communication and immunomodulation. Production of EVs is often increased during viral infections and their role in antiviral immunity is an emerging area of study (32,33). The EVs from human immunodeficiency virus (HIV), Epstein-Barr virus (EBV), and herpes simplex virus 1 (HSV-1) infected cells can have immunomodulatory effects on their hosts through delivery of viral miRNAs or protein products (34–36). Kaposi’s Sarcoma-associated herpes virus (KSHV)-infected cells produce EVs with mitochondrial DNA on their surface that can activate the cGAS-STING pathway (37). Additionally, EVs from U937 cells can deliver poly IC (a synthetic dsRNA) to synovial fibroblasts and exert antiapoptotic effects in a rheumatoid arthritis model (17). Here, we showed that both virus-derived (intracellular) and *in vitro* synthesized long dsRNA (extracellular) can be packaged into EVs, as determined using two different EV isolation methods, and that EVs containing LdsRNA can provide protection against HCoV-229E infection in HEL-299 cells.

A key finding from this work is that long dsRNA, whether it be virus derived or in vitro transcribed, can be packaged into EVs. These LdsRNA-EVs can be extracted using two common EV isolation methods, differential ultracentrifugation and polymer precipitation using a commercially available reagent (ExoQuick-TC^TM^). Both were found to be effective at isolating EVs with detectable LdsRNA. For this study, the ultracentrifugation isolation method was used for the remaining experiments as it provided cleaner and more consistent EV preparations.

An interesting side observation from this study was understanding the stability of intracellular LdsRNA. After 16h treatment, the *in vitro* transcribed dsRNA present in cells was shorter than the original 700 bp dsRNA (HCoV-229E, N), with the majority of the dsRNA being between 150 and 300 bp, while the LdsRNA present in EVs was even shorter, averaging closer to 150 bp (Figure 1B). The decrease in length from intact LdsRNA to intracellular LdsRNA is likely due to degradation within the cell during the 16 hours between treatment and collection. The reduction in size between the cell LdsRNA fraction and the EV fraction initially suggested a size restriction for EV containment. However, this is not true, as VSV-GFP-derived LdsRNA at very long lengths (1000s of bp) was found in the EV fraction (Figure 1D). Thus the reduced size bias seen in Figure 1B dsRNA-EV may be due to low amounts of LdsRNA being detected and not an EV size exclusion phenomenon. The instability in intracellular LdsRNA was observed again with VSV-GFP, where detectable LdsRNA production was consistently observed at 8 hpi (Figure 1D) but less so at 16 and 24 hpi (data not shown). The variability in LdsRNA detection at later time points is likely due to the extensive cell death occurring at 16 and 24h pi. Eight hours is a similar timeframe for LdsRNA detection to what has been observed in other studies using VSV, in which LdsRNA was detected at 6 and 9 hpi (2,25).

Characterization by TEM and Western blot showed that the isolated EV pellets were indeed EVs and that they contained common EV protein markers HSP70 and EMMPRIN (CD147) (29,30). Additionally, histone H3 was detected in both the cell and EV pellets. While histones are not frequently used as EV protein markers, they have been identified in exosomes and microvesicles, particularly during the stress response (31,38,39). The elevated levels of HSP70 and Histone H3 in dsRNA treated cells suggest the dsRNA treatment was triggering a stress response in these cells (Figure 2B). The isolated EVs were roughly 100 nm in diameter, which places them in the small EV (SEV) category (27). Due to their small size and morphology, it’s possible that these SEVs are exosomes, however we did not enrich for any particular subcategory of EV in our extraction, therefore we likely have a mixed population of SEVs (40). SEVs, including exosomes, are known to carry a variety of nucleic acids including mRNA, miRNA, siRNA, circRNA and DNA (reviewed by 43,44). Mitochondrial dsRNA has been found in exosomes from ethanol treated hepatocytes (43), and LdsRNA has been found in the exosomes from neurons following intrathecal morphine injection (44), while poly IC (a synthetic LdsRNA) can be packaged into microvesicles from U937 cells (17).

Prior to antiviral experiments, the ability of EVs to deliver dsRNA to HEL-299 was confirmed through ICC. dsRNA-EVs could deliver long dsRNA to HEL-299 cells and stimulate an antiviral response against HCoV-229E. We saw a significant decrease in the TCID_50_ from cells treated with dsRNA-EVs prior to infection with HCoV-229E (MOI 0.065) indicating that dsRNA-EVs can protect against HCoV-229E infection in HEL-299. No significant reduction in viral titre was observed from the control EVs, therefore it was concluded that EVs alone cannot provide antiviral protection. Additionally, treatment with control or dsRNA-EVs had no significant impact on cell viability in HEL-299. This was unsurprising as EVs are typically found to be non-cytotoxic making them advantageous over other nanocarriers (45–47). The antiviral mechanism induced by dsRNA-containing EVs remains unclear. Both type I interferon signaling and the dsRNA interference (dsRNAi) pathway may have been activated by HCoV-N encoding long dsRNA. Elucidating the relative contribution of these pathways in EV-mediated antiviral responses will be the focus of future studies.

Clearly, cells package LdsRNA into EVs and these EVs have antiviral protection capacity, making them very exciting for future studies. However, we found the dsRNA-EV production methods used in this study to be inefficient, and future studies should explore using cell lines with greater EV secretion such as HEK293 for more scalable production (48). Additionally, there are several cell culturing techniques that could be used to increase EV production such as hypoxia, thermal stress, or media acidification (49–52). Finally, the siRNA profile of EVs from virally infected cells should be studied to determine whether siRNAs matching viral mRNA sequences are present.

Our data support a model in which extracellular vesicles function as vehicles for the intercellular transfer of antiviral LdsRNA, thereby extending LdsRNA-mediated innate immune signaling beyond directly infected cells or cells taking up naked extracellular LdsRNA. This EV-mediated dissemination of LdsRNA may represent a previously unrecognized layer of host defense that primes surrounding tissue during viral infection. Importantly, virus-derived LdsRNA encodes viral sequence that can be processed by Dicer into siRNAs, suggesting that EV-delivered LdsRNA could not only induce type I interferon responses but also preload the RNA-induced silencing complex (RISC) with virus-specific guides in neighboring naïve cells. Such dual activation of IFN and RNAi antiviral pathways would provide a powerful mechanism for anticipatory antiviral defense. Harnessing this natural delivery system could enable the development of scalable, sequence-specific antiviral strategies that leverage endogenous immune pathways rather than synthetic nanocarriers.

## Methods

### Cells and Viruses

U937 cells were obtained from the American Type Culture Collection (ATCC; CRL-1593.2) and maintained in RPMI-1640 (Cytiva) supplemented with 10% fetal bovine serum (FBS, Corning) and 1% penicillin-streptomycin (P/S, Gibco) at 37°C and 5% CO_2_. HEL-299 cells were obtained from the ATCC (CCL-137) and grown in DMEM (Corning) supplemented with 10% FBS and 1% P/S at 37°C and 5% CO_2_. Human coronavirus 229E (HCoV-229E; ATCC; VR-740) was propagated in HEL-299 in DMEM supplemented with 2% FBS at 37°C. At 48 hpi, cells were scraped and resuspended in supernatant. Cells and supernatant were snap frozen in liquid N_2_. Cellular debris was cleared from the media by centrifugation (2700 rcf, 4°C, 10 min) and the supernatant was filtered (0.45µm) and stored at −80°C. The Indiana strain of vesicular stomatitis virus with green fluorescent protein (VSV-GFP) inserted into its genome after the G gene (23) was also used in this study. It was propagated in HEL-299 grown in DMEM supplemented with 2% FBS at 37°C for 48 hrs. Cells and debris were cleared from viral supernatant through centrifugation (2700 rcf, 4°C, 10 min). Viral supernatant was stored at −80°C. For both HCoV-229E and VSV-GFP, propagated virus was quantified by titering on HEL-299 cells to determine the tissue culture infectious dose (TCID_50_).

### Tissue Culture Infectious Dose (TCID_50_)

To quantify HCoV-229E or VSV-GFP, HEL-299 were seeded in 96 well plates at a density of 4.8×10^4^ cells/well. Viral supernatant was then serially diluted from 1.0×10^−1^ to 1.0×10^−8^ in DMEM supplemented with 2% FBS. 100 µL of each dilution was added to cells in replicates of 6 wells per dilution. Cells were examined for cytopathic effects (CPE) after 6 days post infection (dpi) for HCoV-229E or 3 dpi for VSV-GFP, and the Reed and Meunch method was used to calculate the TCID_50_ (53).

### In vitro transcribed (ivt) Long dsRNA Synthesis

A template for LdsRNA synthesis was prepared by cloning complementary DNA (cDNA) encoding 700 nucleotides of the HCoV-229E genome (bases 25992 −26691) into the pGEM^®^-T vector (Promega). The HCoV-229E N gene was amplified through PCR using forward and reverse primers containing the T7 RNA polymerase promoter sequence at their 5’ ends (Table 1) (5). The PCR reaction contained 10 ng of template plasmid, 0.5µM each of forward and reverse primer and 2X GoTaq Colourless Master Mix (Promega) and brought up to 50µL with nuclease free water. The reaction was run in a BioRad T100 Thermal Cycler set to 95°C for 5 min followed by 40 cycles of 95°C for 30 s, 58°C for 30 s, and 72°C for 1 min, followed by a final extension at 72°C for 5 min. The PCR product was purified using the QIAquick PCR Purification Kit (QIAgen). Two µg of purified PCR product was used in each dsRNA synthesis reaction using the MEGAscript RNAi kit (Invitrogen) following manufacturer’s directions including purification steps to remove contaminating ssRNA and dsDNA. The resulting dsRNA concentration was determined using a NanoDrop Lite Spectrophotometer (Thermo Scientific) and purity was assessed through 1% agarose gel electrophoresis stained with 1X RedSafe (iNtRON Biotechnologies).

**Table 1.**
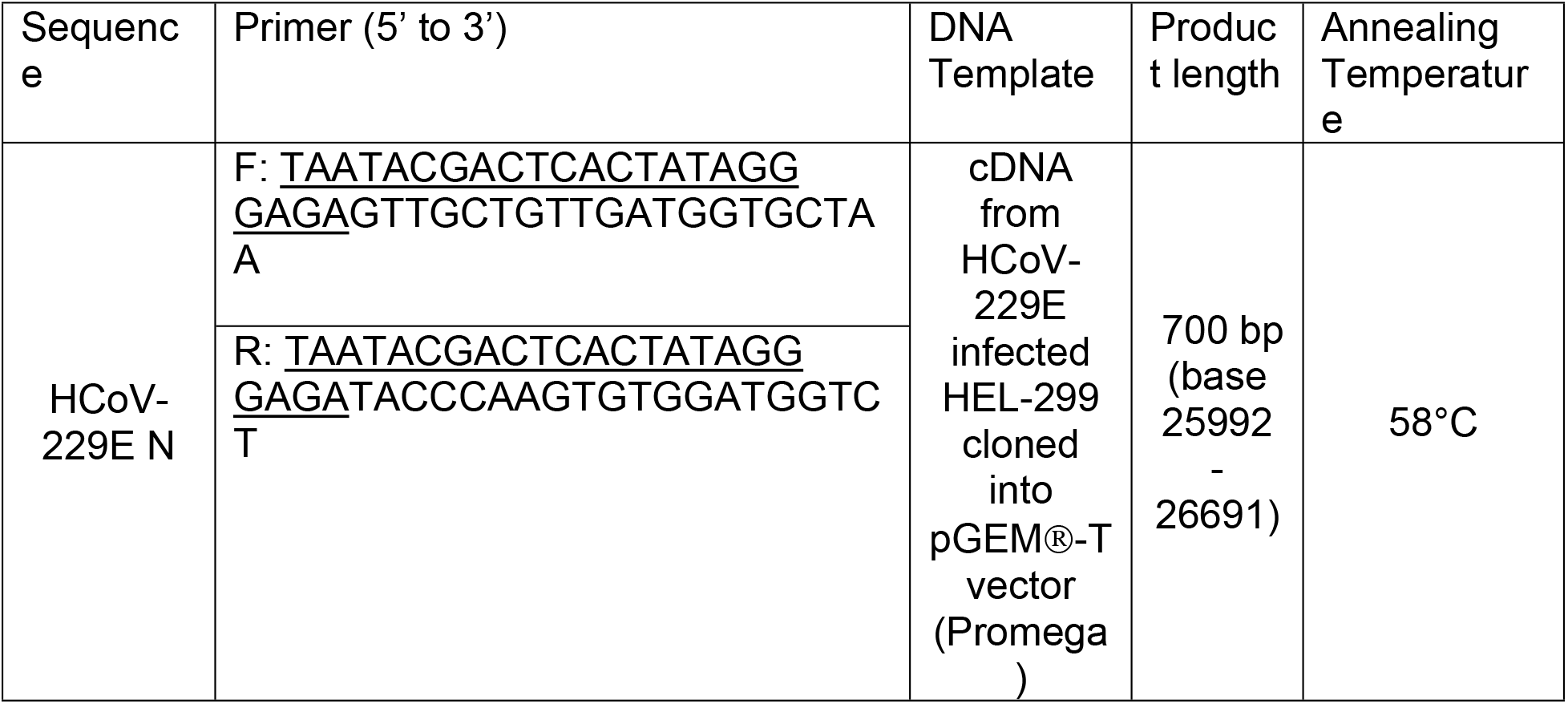
PCR primer sequence for dsRNA synthesis. Underlined nucleotides make up sequence for T7 RNA polymerase promoter.

### LdsRNA treatment and EV isolation by ultracentrifugation

U937 were seeded at 1.0×10^7^ cells/ flask in a 25 cm^2^ tissue culture treated flask and stimulated with 1.5 ng/mL phorbol myristate acetate (PMA). Forty-eight hrs post treatment, cells were washed with DPBS to remove unstimulated cells. One flask was treated with 33.3 µg/mL long dsRNA (HCoV-229E N gene, 700 bp) in basal RPMI-1640 media, while a control flask was given only basal RPMI-1640. Sixteen hrs post treatment, cells were collected by centrifugation (2700 rcf, 15 min; cell fraction) and supernatants were filtered (0.22 µm). The 16h post treatment collection point was based on previous studies (17,54). EVs were pelleted from cleared media (100,000 rcf, 2 hrs, 4°C) (55). The supernatant was then removed and pellets were then washed twice in DPBS (100,000 rcf, 30 min, 4°C; EV fraction). EVs isolated from LdsRNA-treated cells are termed dsRNA-EVs, while EVs isolated from untreated cells are termed control EVs.

### VSV-GFP Infection and EV isolation by exosome precipitation

To determine optimal collection time of EVs following VSV-GFP infection, U937 cells were plated at a density of 1.0×10^7^ cells/mL in a 25 cm^2^ tissue culture flask and stimulated with 1.5 ng/mL PMA. 24 hrs post treatment, cells were washed with DPBS and then infected with VSV-GFP (MOI 1) for 2 hrs in 3 mL basal media. 3 mL of media were added 2 hpi and FBS was added to a concentration of 2%. At 8, 16, and 24 hpi, cells were imaged at 200X magnification using a Nikon Ti2 microscope with FITC and brightfield channels to assess production of GFP by VSV-GFP and resulting cytopathic effect (CPE).

To detect VSV-GFP dsRNA in EVs, cells were plated and infected with VSV-GFP (MOI 1) as described above. At 8 hpi cells were scraped from the flask and pelleted at 600 rcf (cell fraction). Cell debris was cleared from media by centrifugation at 1400 rcf for 10 min at room temperature, followed by syringe filtration with a 0.22µm filter. Cleared media was mixed with 1 mL ExoQuick-TC™ (Systems Bioscience) and left to precipitate EVs overnight at 4°C. EVs were pelleted from media by centrifugation at 2700 rcf for 30 min at room temperature (EV fraction). RNA was extracted from cell and EV pellets as described above.

### RNAse III Treatment of EVs from VSV-GFP infected U937

U937 were plated as described above and infected with VSV-GFP (MOI 1). At 8 hpi, cells were collected and media was cleared of cells and debris as described above. One µg LdsRNA (HCoV-229E N, 700 bp) was spiked into cleared media prior to the addition of 500 µL ExoQuick-TC™. EVs were pelleted (2700 rcf, 30 min) after overnight precipitation at 4°C. EV pellets were resupended in 88 µL nuclease-free water and mixed with 2U RNase III (Ambion) and 10 µL 10X RNase III Reaction Buffer (Ambion) and incubated at 37°C for 1 hr. Samples were mixed with 500 µL TRIzol™ and RNA was extracted following manufacturers directions for immunoblotting as described below.

### Transmission Electron Microscopy

EV pellets isolated by ultracentrifugation were resuspended in 100µL of phosphate buffered saline (PBS) supplemented with 25 mM HEPES and 0.2% w/v BSA (pH 7.4) (56). Pellets were frozen at −80°C until imaging by transmission electron microscopy (TEM) at the University of Guelph Advanced Analysis Centre. Prior to imaging, EVs were bound to copper grids for 2 min and washed three times with 20 mM HEPES buffer. EVs were then stained with 2% uranyl acetate for 10 seconds and imaged at 50,000X magnification.

### Western Blot

Cell and EV pellets were resuspended in 500 µL RIPA buffer (Sigma-Aldrich) and incubated on ice for 30 minutes. Samples were centrifuged at 13,000 rpm for 13 min and protein contents of supernatants were quantified using the Pierce BCA Protein Assay (ThermoFisher). 12 µg of protein was separated by 10% SDS-Poly acrylamide gel electrophoresis (PAGE). Proteins were transferred onto a nitrocellulose membrane via the Trans-Blot^®^ Turbo™ Transfer System (Bio-Rad 1704150). A 0.1% Ponceau S. stain was applied for 1 min with rocking followed by 3 rinses in MilliQ water (5 min each). The Ponceau S. stain was imaged using visual light to ensure equal protein loading and the stain was then removed with Tris-buffered saline supplemented with 0.1 % Tween 20 (TBS-T). The membrane was blocked in 5% skim milk powder dissolved in TBS-T for 1 hr with rocking. The blocking solution was then removed and the membrane was rinsed 3 times (5 min each) with TBS-T. Membranes were then probed with anti-HSP70 (Sigma-Aldrich), anti-Histone H3 (Cell signalling), or anti-EMMPRIN (Millipore Sigma) primary antibodies diluted 1:2000 in TBS-T with 5% w/v BSA overnight at 4°C with rocking. Primary antibody solutions were then removed, the membrane was rinsed 3 times with TBS-T and the appropriate secondary antibody [Goat anti-mouse HRP (Biotium) for HSP70 and Goat anti-rabbit HRP (Bio-Rad) for EMMPRIN and Histone H3] diluted 1:4000 in blocking solution was applied for 1 hr with rocking. The membranes were then rinsed twice with TBS-T and once with TBS prior to incubation in Clarity Western ECL Substrate (Bio-Rad) for 5 min in the dark. Blots were imaged via chemiluminescence using the Azure Biosystems c280 imager.

### dsRNA immunoblots

EV and cell pellets were resuspended in RIPA buffer (Sigma-Aldrich) and incubated on ice for 30 min to allow lysis. Two volumes of 100% isopropanol and 8µg RNA-grade glycogen were added and incubated at −80°C for 6 hrs to allow RNA precipitation. RNA was pelleted at 21130 rcf for 15 min at 4°C and washed with 75% v/v ethanol (21130 rcf, 5 min, 4°C). Each pellet was dried for 5 min at room temperature, resuspended in 20 µL nuclease-free water and incubated at 55°C for 15 min. An immunoblot with the J2 anti-dsRNA antibody was performed on RNA extracted from cell and EV pellets as described in Poynter and DeWitte-Orr (2017) (57). Briefly, RNA samples (2µg) were separated by 10% PAGE (140V, 115 min). The RNA was then transferred to a positively charged nylon membrane (Roche; 25 V, 90 min) using the Trans-Blot^®^ Turbo™ Transfer System (Bio-Rad 1704150). The RNA was crosslinked to the membrane with UV light for 20 min using the auto-crosslink function of the Stratalinker® 1800 (Stratagene). The membrane was blocked in 5% skim milk powder in TBS-T for 1 hr with rocking. The blocking solution was then removed and the membrane was rinsed 3 times (5 min each) with TBS-T. The membrane was incubated in the J2 anti-dsRNA antibody (Novus) diluted 1:1000 in TBS-T overnight with rocking followed by rinsing 3 times (5 min each) with TBS-T and an incubation in goat-anti mouse secondary antibody (Biotium) diluted 1:2000 in TBS-T for 1 hr with rocking. The membrane was then washed twice with TBS-T and once with TBS (5 min each) and then incubated in Clarity Western ECL Substrate for 5 min (Bio-Rad). Blots were imaged with the Azure Biosystems c280 imager using chemiluminescence with cumulative autoexposure.

### Immunocytochemistry

HEL-299 cells were seeded at a density of 9.25×10^4^ cells/chamber in an 8-well µ-slide (ibidi) in DMEM medium (10% FBS, 1% P/S) and allowed to attach overnight. Cells were treated for 16 hrs with dsRNA-EV pellets or control-EV pellets derived from T25 flasks, resuspended in 40µL basal DMEM. A positive control well was treated with 5 µg N dsRNA, and untreated control wells received basal media only. After 16hrs, treatments were removed and cells were washed three times with PBS (1 min each). Cells were fixed to the slide flask with 10% neutral buffered formalin for 10 min followed by three washes with PBS. Cells were permeabilized with 0.5% Triton-X in PBS solution for 15 min followed by three rinses in PBS. Cells were blocked overnight in PBS with 3% w/v BSA, 3% v/v goat serum and 0.02% v/v Tween-20 at 4°C in a humidified chamber. Cells were then rinsed three times with PBS and treated with the J2 primary antibody diluted 1:200 in blocking buffer for 1 hr. Cells were washed three times in PBS and then treated with a TRITC conjugated goat anti-mouse secondary antibody (Invitrogen) for 1 hr. Cells were washed three times with PBS and nuclei were counterstained with 5µg/mL DAPI in PBS for 5 min in the dark followed by two additional rinses in PBS and one in MilliQ water. Cells were imaged at 200X magnification with a Nikon Ti2 Eclipse microscope using epifluorescence.

### EV Cytotoxicity (Alamar Blue)

HEL-299 were seeded in a 48 well plate at a density of 9.25×10^4^ cells/well and allowed to attach overnight. dsRNA-EV and control EVs were extracted from ivt-dsRNA treated and untreated control U937 cell culture by ultracentrifugation as described above. EV pellets were resuspended in 200 µL basal DMEM. HEL-299 were treated with resuspended EVs for 16 hr. Cells were rinsed with DPBS and treated with 5% alamarBlue (Invitrogen), an indicator of cellular metabolism, diluted in DPBS for 1 hr at 37°C with 5% CO_2_. Fluorescence at 560/590 was measured in a Varioskan LUX microplate reader (ThermoFisher).

### Antiviral Assay

HEL-299 cells were seeded at a density of 9.25×10^4^ cells/well in a 48 well tissue culture plate in DMEM medium (10% FBS, 1% P/S) and allowed to attach overnight. Cells were treated with dsRNA-EV or control-EV pellets isolated by ultracentrifugation from ivt-dsRNA and untreated control U937 cells in T25 flasks, resuspended in 200 µL basal DMEM. At 16 hours post treatment, the media was removed, and cells were washed with DPBS. Cells were then infected with HCoV-229E (MOI 0.065) in 300 µL basal DMEM. Two hours post infection, the media was removed and replaced with DMEM supplemented with 2% FBS. To measure resulting viral titres, the supernatant was collected 48 hpi and stored at −80°C. The viral titres were measured via tissue culture infectious dose (TCID_50_) assays on HEL-299 cells as described above.

### Statistics

All graphs and statistical analysis were prepared in GraphPad Prism 10. Data are shown as the mean average with error bars representing the standard error of the mean (SEM). Statistical analysis was performed using a Kruskal-Wallis test with a Dunnett’s post hoc test where a p-value < 0.05 was considered significant. HCoV-229E viral titres were log transformed to normalize data. Outliers were determined using the Robust regression and Outlier removal (ROUT) method with Q= 1%.

## Acknowledgements

The authors wish to thank Dr. Erin Anderson from the University of Guelph Advanced Analysis Centre for imaging the EVs through TEM. The authors also wish to thank Dr. Shayne Oberhoffner, Sarah Au, and Dr. Sarah Poynter for their guidance and contributions to experimental design.

